# Assigning a role for chemosensory signal transduction in *Campylobacter jejuni* biofilms using a combined omics approach

**DOI:** 10.1101/862151

**Authors:** Greg Tram, William P. Klare, Joel A. Cain, Basem Mourad, Stuart J. Cordwell, Victoria Korolik, Christopher J. Day

**Author notes:** Corresponding Authors (VK), (CD). These authors contributed equally to this work (Greg Tram, William P. Klare). These authors contributed equally to this work (Victoria Korolik, Christopher J. Day).

## Abstract

Biofilms serve as a protective mechanism for bacteria to cope with environmental stress. Whilst ordinarily a fastidious organism, it has been long suggested that *C. jejuni* is able to utilise this mode of growth as a way to transmit infection from the avian host to humans. Herein, we undertook a combinatorial approach to examine differential expression of *C. jejuni* genes and protein abundance during biofilm formation. RNA sequencing and proteomics *via* quantitative liquid chromatography – tandem mass spectrometry (LC-MS/MS) revealed biofilm growth induced a substantial rearrangement of the *C. jejuni* transcriptome and proteome, with ∼600 genes differentially expressed in biofilms compared to planktonic cells. Biofilm-induced genes / proteins were those involved in iron metabolism and acquisition, cell division, glycan production and attachment, while those repressed were associated with metabolism, amino acid uptake and utilisation, and large tracts of the chemotaxis pathway.

We further examined the role of chemotaxis in *C. jejuni* biofilm formation by assessing the behaviour of isogenic strains with deletions of the *cheV* and *cheW* genes. Both Δ*cheV* and Δ*cheW* exhibited a significant decrease in directed motility when compared to wild-type *C. jejuni*. Both mutants also demonstrated an increase in autoagglutination ability and increased biofilm formation. A subtle difference was also observed between the phenotypes of Δ*cheV* and Δ*cheW* mutants, both in motility and biofilm formation. This suggests roles for the CheV and CheW signal transduction proteins and may present signal transduction as a potential method for modulating *C. jejuni* biofilm formation.

**Author summary:** *Campylobacter jejuni* is a gastroenteric bacterium that is responsible for most cases of bacterial food poisoning in the developed world. The organism commonly resides in avian reservoirs and is passed to humans through contaminated poultry and animal products. Ordinarily, *C. jejuni* requires a strict set of conditions in order to survive and cause infections in humans. Biofilms are a method of bacterial growth that may provide shielding from harsh environments and provide an important link between reservoirs and human hosts. In this study, we have utilised a multi-platform approach to compare gene expression and protein abundance in planktonic *C. jejuni* cells and those growing as a biofilm. We subsequently focused on the chemosensory system of *C. jejuni* and demonstrated that signal transduction proteins play a role in biofilm formation. Our work has provided a broad profile of which genes are important to *C. jejuni* biofilms and that the chemosensory pathway has an influence on biofilm formation.

## Introduction

*Campylobacter jejuni* is a Gram-negative bacterium responsible for the majority of bacterial gastroenteritis cases in developed nations. *C. jejuni* is a common commensal organism of avians and is transmitted to humans through contaminated food products or water. Once ingested, *C. jejuni* migrates through the gastrointestinal tract, colonising, and infecting cells of the intestine [1]. *C. jejuni* infections have also been implicated in the development of post infection complications such as the autoimmune neuro-degenerative conditions, Guillain-Barré and Miller-Fisher syndromes [2].

*C. jejuni* biofilms have been suggested to play a role in the transmission of infection between chickens and human hosts [3]. Exacting and microaerophilic, *C. jejuni* may utilize biofilms as a method of surviving nutrient deficient and oxygenated environments, such as food preparation and storage [4], in order to remain viable. To date, a number of factors have been shown to play a role in *C. jejuni* biofilm formation. A large number of protein families have been implicated, including glycosylation pathways [5], DPS (DNA-binding protein from starved cells) proteins [6], and post-transcriptional regulators [7]. An interesting aspect of *C. jejuni* biofilm formation is the impact of motility and chemotaxis gene exression [8,9].

Chemotaxis in *C. jejuni* is a two-component signal transduction system where a change in conformation of the chemoreceptors, known as transducer-like proteins (Tlps), directs autophosphorylation of a histidine kinase, CheA. CheA phosphorylates CheY, which acts as a cytoplasmic Response Regulator (RR) that migrates to the flagella motor where it can influence changes in flagellar rotation. The phosphorylation of CheAY also serves to activate CheB, a methyltransferase that demethylates the signalling domain of Tlps upon the binding of chemorepellents. *C. jejuni* possesses an additional scaffolding protein, CheV. CheV is comprised of a CheW-like domain and an additional CheY-like RR domain at the *C*-terminus. The CheW-like domain of CheV has been shown to play a similar role to CheW and the two share conserved sequence [10,11]. Currently, the function of the RR domain of CheV is unknown, although there is evidence it may be involved in sensory adaptation probably because CheB of *C. jejuni* lacks an RR domain present in CheB proteins of other bacteria [12]. A number of *C. jejuni* chemoreceptors have been shown to have a higher affinity for either CheV or CheW in *C. jejuni*. The Tlp2, Tlp3 (CcmL), Tlp4 paralogues and Tlp11 (CcrG) chemoreceptor signalling domains have been shown to interact non-preferentially with either CheV or CheW proteins [13-15], whereas the Tlp1 chemoreceptor has demonstrated strong interactions with only the CheV homologue [16].

Elements of this chemotaxis pathway have been shown to play a significant role in the formation of biofilm in *C. jejuni.* General motility is required, as deletion of components of the flagella complex results in a loss of biofilm forming ability [17]. Interestingly, chemoreceptors appear to influence both increases and decreases in biofilm formation. Deletions in Tlp8, for example, result in decreased biofilm production, whereas deletions in Tlp3 increase biofilm levels. This may be due to the effect of chemoreceptors on motility, as the Tlp8 mutant demonstrates increased motility whilst the directed motility of a Tlp3 mutant is diminished due to its “tumble” phenotype bias [8,14].

In this study, we utilize a comparative omics approach to reveal genes and proteins that are associated with *C. jejuni* biofilm growth. Analysis of the transcriptome and proteome demonstrated distinct variations between planktonic and biofilm states and revealed key roles for a number of families of proteins including those involved in chemotaxis. We further investigated the relationship between the chemotaxis pathway and *C. jejuni* biofilms, by specifically examining the role of the chemotaxis proteins CheV and CheW in *C. jejuni* biofilm formation.

## Results

### RNA transcription during planktonic and biofilm growth states

Detailed methods and accession numbers for RNA-seq datasets can be found in the corresponding Microbial Resource Announcement [18]. RNA-seq analysis identified 1571 genes equating to ∼96.8% coverage of the *C. jejuni* NCTC 11168 genome (S1 Table). 789 genes were considered as differentially expressed (DE) using a fold change cut-off of Log_2_ <-1 and >+1 (equivalent to a +/-2-fold change), with 427 up-regulated and 362 down-regulated transcripts in biofilm conditions (S2 Table). Increasing the stringency by including only those DE genes that met the false discovery rate (FDR) threshold of <0.01 removed 169 genes from further analysis. Therefore, a final total of 620 genes were considered DE at the +/-2-fold level of regulation, with 318 up- and 302 down-regulated genes (Fig. 1, S2 Table).

**Fig 1.**
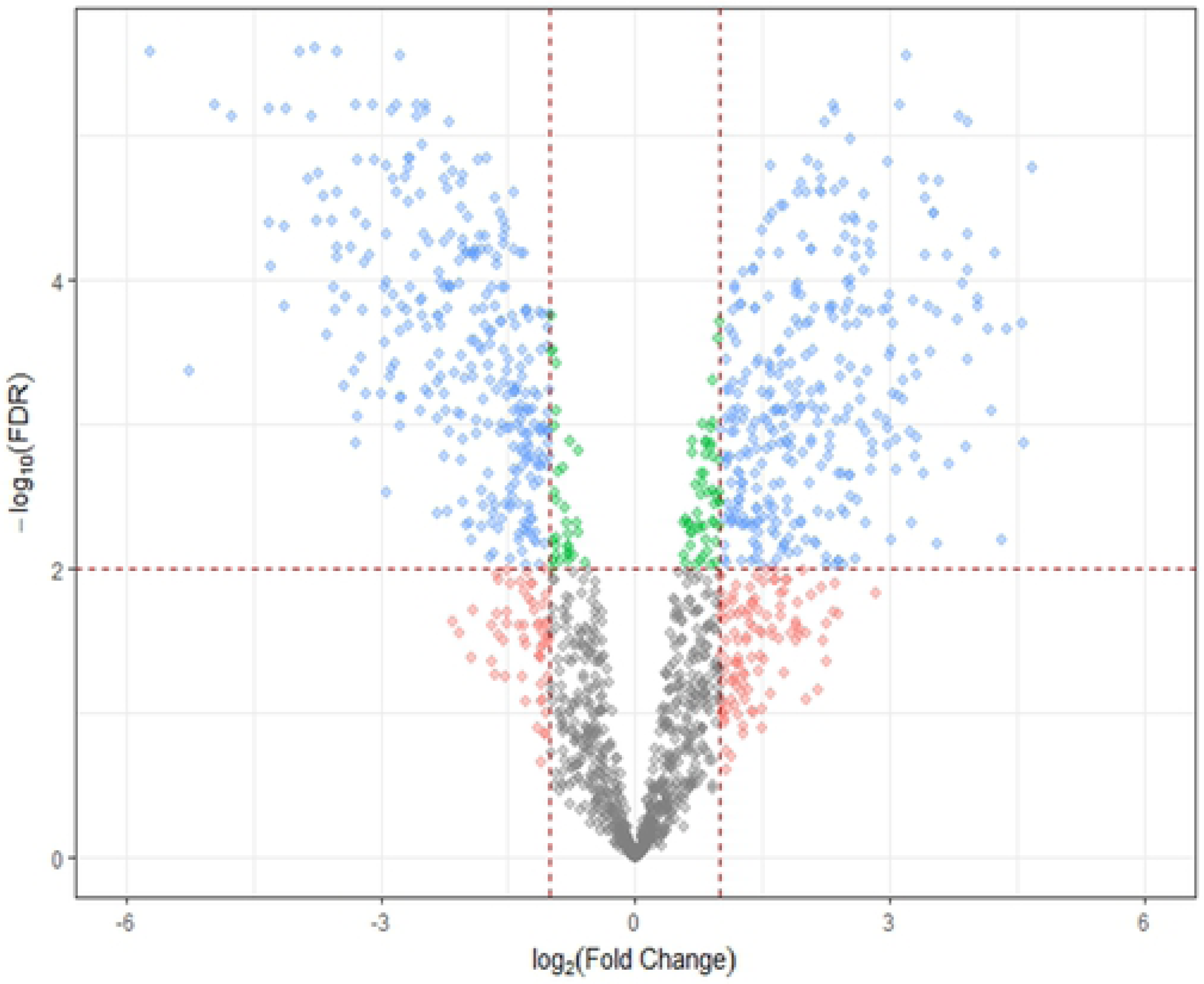
Volcano plot of *C. jejuni* NCTC 11168 gene expression during biofilm growth. Log2 (Fold Change) significance cut-off was +/-1, with threshold for significance set at -log10 (FDR) <2. A total of 1571 genes were identified (96.7% coverage) and 620 were significantly differentially expressed (blue shaded data points).

These DE genes were next subjected to functional cluster analysis using STRINGdb. DE genes that were up-regulated in *C. jejuni* NCTC 11168 biofilm growth (S1a Fig.) were strongly associated with several functionally related clusters; including iron limitation and iron acquisition (enterochelin uptake, *ceuBCD;* and haemin uptake, *chuABCD; cfbpBC, cfrA* and several unique genes e.g. *cj1384c* and *cj0730*), ribosomal proteins and translation, amino acid biosynthesis and metabolism also related to signal molecule production and biofilm formation (*metABEF*; *trpABD, hisABF1*), cell division and peptidoglycan (*pbpAC, murG, mreB*) and production and attachment of *N*-glycans (e.g. *pglABCDIG*). Several smaller clusters were broadly related to transport of nutrients and small molecules (e.g. molybdenum, *modABC*; phosphate, *pstABC*), branched chain amino acid transport (*livFGHKM*) and co-location in operons with no currently assigned function (e.g. *cj0727-cj0732*; *cj1658-cj1663*; *cj1581c-cj1583c*). We also examined a subset of these DE genes that exhibited the greatest magnitude of change (>+5-fold; S1b Figs.). This emphasised the changes associated with iron acquisition and iron-regulated genes; as well as molybdenum and phosphate acquisition, and clusters of genes with no known function (e.g. *cj1660-cj1662*; *cj1581c-cj1583c*). Down-regulated DE genes (S1c Fig.) clustered into fewer functional groups, however major clusters were associated with chemotaxis and transducer-like proteins (*cheBRWY* and *cj0019c, cj0144, cj1506c, cj1564*; as well as *cj0448c* [homologous to biofilm dispersion protein *bdlA*]), amino acid uptake, utilization (*sdaC, dcuA, aspAB*) and metabolism (*sdhABC*), nitrate reductase (*napABDGHL*) and motility (*flgABCHM, fliEL, flaG*). Smaller clusters were again related to specific operons (e.g. *cj1430c-cj1436c, cj0770c-cj0772c*; *cj0920c-cj0922c* [encoding *peb1A* and *peb1C*] and *cj0552-cj0554*). Similar to the analysis performed for up-regulated genes, we next specifically looked at genes with large magnitude down-regulated fold changes (<-5-fold; S1d Fig.), which further highlighted changes associated with motility, nutrient uptake and metabolism, and related genes of no known function.

### LC-MS/MS proteomics of *C. jejuni* NCTC 11168 biofilm growth

Label-based proteomics analysis by trypsin digest and LC-MS/MS was performed on biological triplicates and 1278 proteins (∼78.7% of the 1623 *C. jejuni* NCTC 11168 genes in the Uniprot database) were identified in at least 1 biological replicate (S3 Table) by a minimum of 2 or more unique peptides. 11 proteins were identified where the corresponding transcripts were not detected by RNA-seq (Cj0185c, Cj0202c, Cj0455c, Cj0483, Cj0679, Cj0874c, Cj0876c, Cj0951c-Cj0952c, Cj1392-Cj1393). Log_2_ fold changes were converted to *n*-fold change for further analysis. Similar to the data observed by transcriptomics, biofilm growth resulted in a major remodelling of the *C. jejuni* proteome and 431 proteins were considered differentially abundant (DA) at a fold-change cutoff of +/-1.5-fold (S4 Table) with 220 present at increased abundance and 211 present at reduced abundance in *C. jejuni* biofilm growth. As for the DE gene expression analysis, we next subjected the DA proteins to functional cluster analysis using STRINGdb. The DA proteins present at increased biofilm abundance displayed major clusters associated with ribosomal proteins (translation), protein *N*-glycosylation and intracellular metabolic processes, including methionine metabolism (MetABEF; S2 Fig.). Another cluster was associated with antimicrobial resistance (predominantly CmeABC). DA proteins present at reduced levels in biofilm growth clustered into several major groups (S2b Fig.), including ribosomal proteins, motility and chemotaxis proteins (FlgD, FliN, CheRY), proteins involved in respiration that allow the use of alternative electron acceptors under low oxygen conditions (e.g. nitrate reductase NapABDGL, formate and fumarate dehydrogenases [FdhABC; SdhABC] and Cj0264c-Cj0265c), nutrient acquisition and utilization (AspAB, AnsA, DcuA), a group of proteins involved in glycan biosynthesis (Cj1427c-Cj1438c), and a small cluster of functionally unknown proteins (Cj0411-Cj0413).

We next integrated the omics data to identify genes and their products that were closely associated with this model of biofilm growth in *C. jejuni*. The RNA-seq and proteomics data were aligned by fold-change and statistical analysis (Fig 2a). The data showed that transcript and proteomics data correlated well (Spearman coefficient 0.767 with *p*<0.001), although fold changes were generally considerably higher for the transcriptomics data (−52.33-to +25.45-fold) compared with the proteomics data (−9.38-to +14.42-fold). We also observed that larger RNA-seq-derived fold changes, particularly those that were up-regulated DE genes, mainly correlated with proteins that could not be detected by LC-MS/MS (of 48 genes up-regulated >8-fold only 11 could be identified at the protein level), most likely indicating that these transcripts (and proteins) are of lower starting abundances. Examination of the entire aligned dataset (transcripts and proteins both identified) using heat mapping software also showed that the two datasets typically correlated (as measured by direction of any expression / abundance change), although some specific outliers could be observed (Fig. 2b).

**Fig 2.**
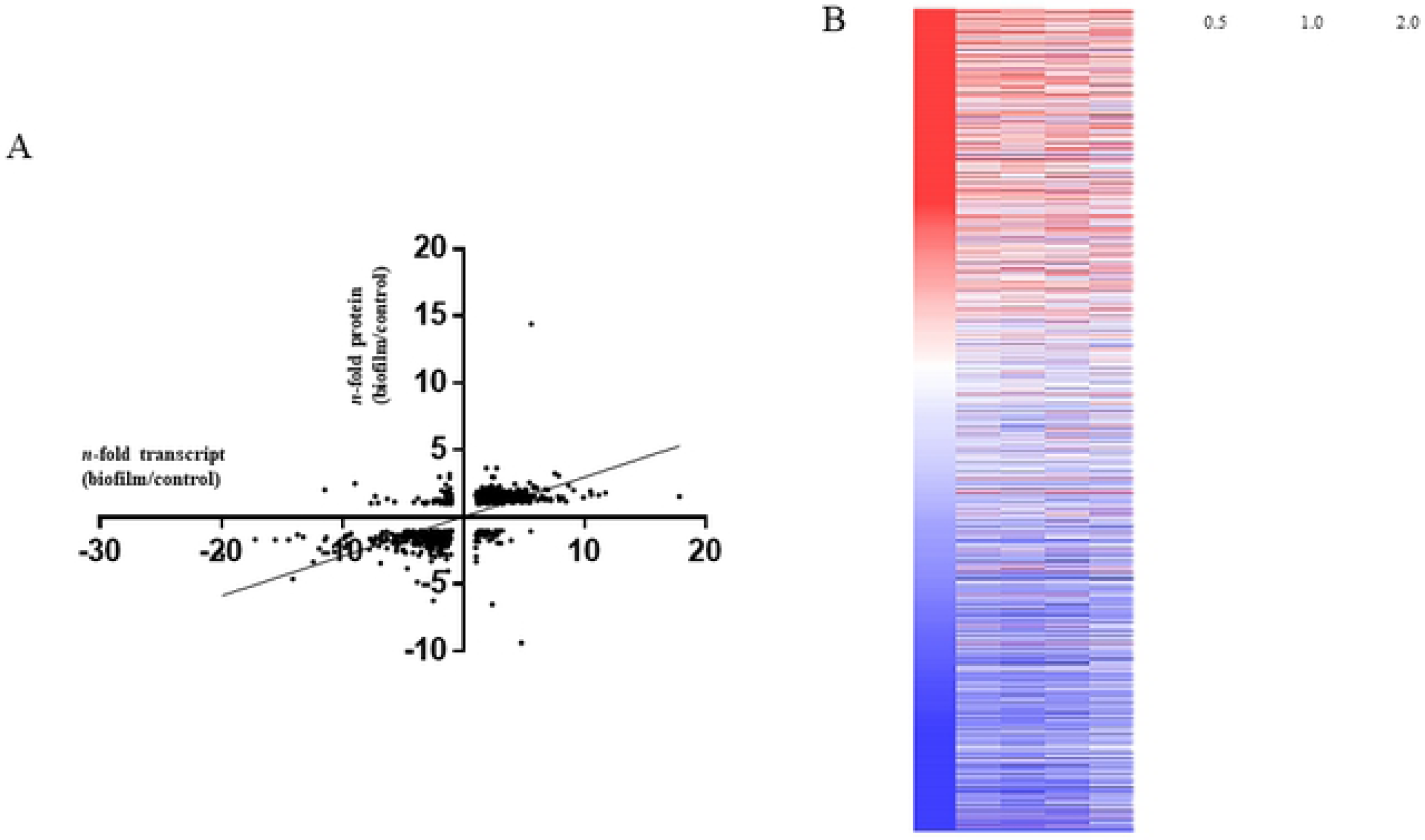
Comparisons of transcriptomics and proteomics. (A) Correlation plot of gene expression (x-axis) versus proteomics data y-axis). Spearman correlation coefficient = 0.767; (B) Heat map of gene expression (left column) and proteomics (left to right; mean; biological triplicate data) data, ranked according to fold change as determined by RNA-seq gene expression.

We considered such ‘outliers’ to be defined as a gene/protein with an opposite and significant DE/DA ratio (e.g. a significantly down-regulated gene showing a significantly up-regulated protein abundance). Only 11 genes/proteins were outliers; 7 with elevated gene transcript and significantly reduced protein abundance (*rpmG, fdhU, cj0299, cj0908, cj1169c, ybeY* and *cj0648*), and 4 with reduced transcript level and increased protein abundance (*dps, ciaI, ftn* and *cj1053c*) (S5 Table).

### Biofilm formation

Since our integrated omics analysis strongly indicated repression of genes / proteins involved in chemosensing and chemotaxis (Fig. 3A and Fig. 3B), we next examined the role of these functions in *C. jejuni* biofilms. Quantitative comparison of biofilm formed by wild-type *C. jejuni* NCTC 11168 O and *ΔcheV* and *ΔcheW* chemotaxis-deficient mutants, demonstrated that the *cheV*-deficient isogenic mutant strain had an approximately five-fold increase in biofilm formation, whilst the *cheW*-deficient mutant showed an ∼three-fold increase in the ability to form biofilms (Fig. 3C).

**Fig 3.**
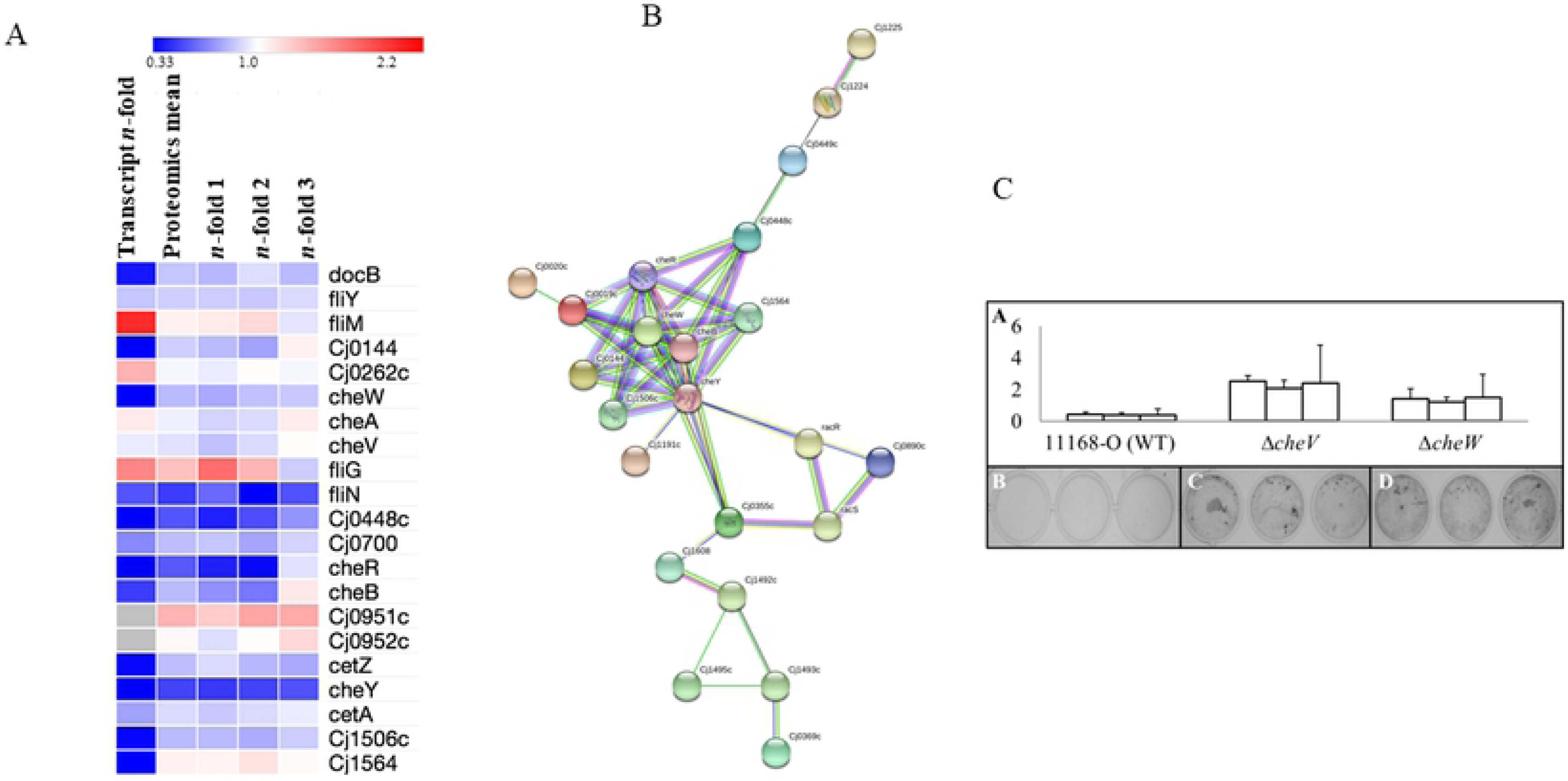
Association of chemotaxis and chemosensory genes / proteins with biofilm formation. (A) Heat map showing gene expression and relative protein abundance of chemosensory and chemotaxis-associated genes during biofilm growth; (B) STRINGdb cluster of enriched chemotaxis-associated genes; (C) Dissolution of stained biofilm showing increased levels of biofilm formation, measured by absorbance at 600nm, in CheV and CheW deficient mutants, (D,E,F) images of stained biofilm in 24-well plates show this same increase qualitatively.

### Motility and autoagglutination of che mutants

Motility assays were conducted in order to compare the ability of wild-type, *ΔcheV* and *ΔcheW C. jejuni* to migrate through a solid support. Wild-type *C. jejuni* NCTC 11168 O showed an average migration of 28 mm from the inoculation site (Fig. 4A and 4B). Both *ΔcheV* (4.5 mm migration) and *ΔcheW* (6 mm migration) mutants showed significantly diminished motility when compared to the wild-type strain (*p* value of 0.033; Fig. 4A, 4C and 4D).

**Fig 4.**
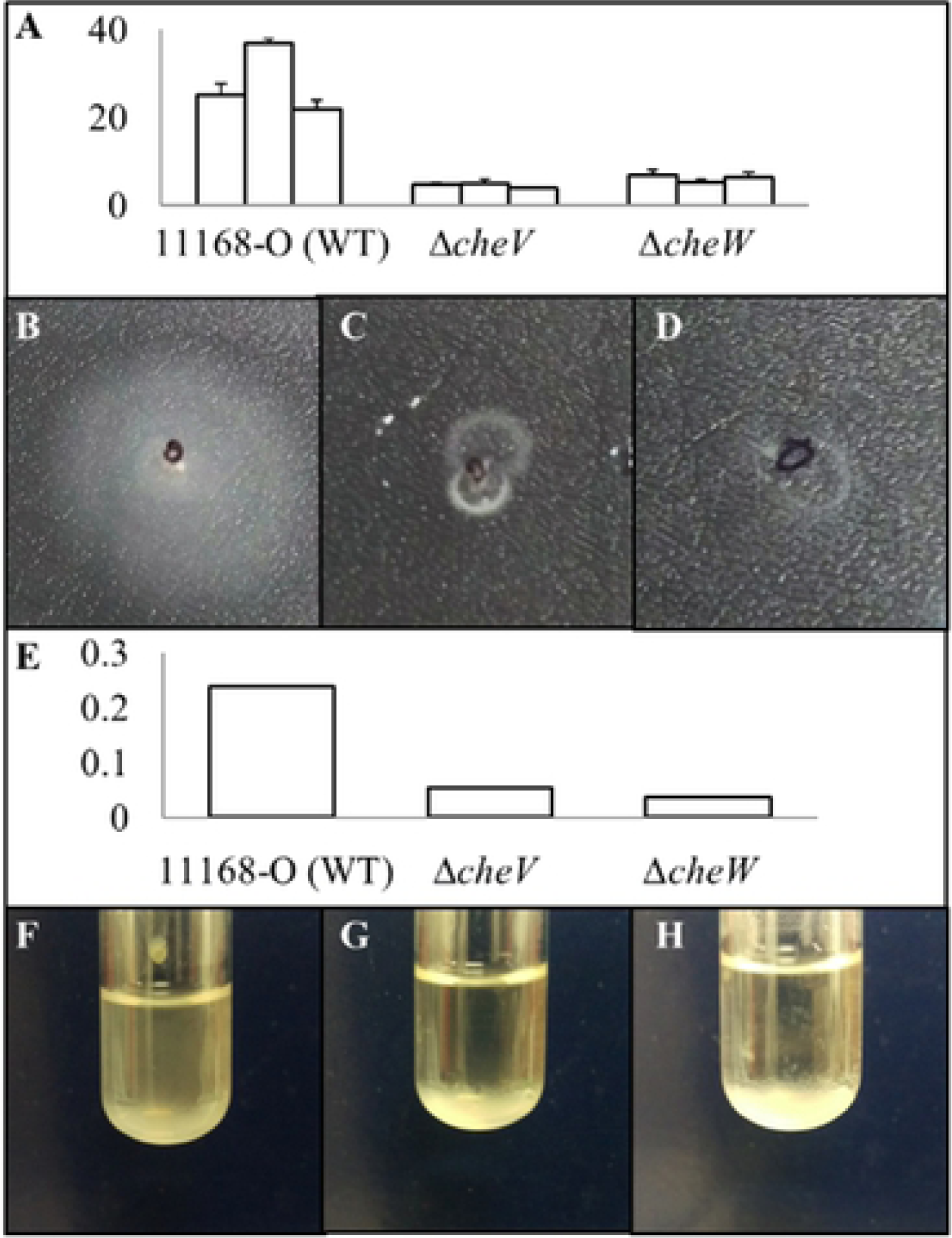
Analysis of motility and autoagglutination activity. (A) Migration (mm) of bacteria from the inoculation site for WT, *ΔcheV* and *ΔcheW* isogenic mutants in triplicate; (B,C,D) Images of the formed halos demonstrating decreased swarming motility by WT, *ΔcheV* and *ΔcheW* mutants; (E) Spectrophotomer readings of the upper phase of cell suspension following liquid culture demonstrate a lower absorbance in *ΔcheV* and *ΔcheW* strains when compared to WT, indicating an increased tendency for cells to autoagglutinate, (F,G,H) which was also evident when culture tubes were inspected.

Autoagglutination assays were carried out to determine the impact of diminished motility on intercellular adherence. Cells with a higher tendency to adhere to neighbouring cells cause increased settling and a lower absorbance of the upper phase. Both *ΔcheV* and *ΔcheW* mutants (Fig. 4G and 4H) showed a marked increase in clustering when compared to the wild-type control (Fig. 4F). This increase in autoagglutination was confirmed by assessing the absorbance of the upper phase of media in the culture tubes. A marked decrease was noted for both mutants (agglutination assays performed in triplicate), suggesting a large increase in autoagglutination (Fig. 4E).

### Time-lapse microscopy

Time-lapse microscopy was conducted to highlight differences in overall biofilm formation in *C. jejuni* isogenic strains (S3 Fig.). Both *ΔcheV* and *ΔcheW* isogenic mutants exhibited a higher tendency to aggregate into cell clusters than wild-type *C. jejuni.* This is particularly evident during the interim stages of biofilm development such as seen at 15-20 hours. Wild-type *C. jejuni* showed a progressive spread of biofilm formation into a confluent monolayer, whereas *ΔcheV* and *ΔcheW* mutants appeared to congregate more readily into large microcolonies. In addition to the increased aggregates, a much larger biomass was evident in the *ΔcheV* and *ΔcheW* mutants when compared to the wild-type.

## Discussion

Combining the use of RNA sequencing and proteomics enabled a global understanding of *C. jejuni* biofilm growth and demonstrated that significant remodelling of the transcriptome and proteome occurs in this lifestyle compared with planktonic culture at 42°C. We identified many genes encoding protein families that were differentially expressed in biofilms including those involved with glycosylation, metabolism and chemotaxis, as well as a large number of currently functionally poorly characterised gene clusters.

Generally, chemotaxis has long been established to play an important role in biofilm formation in *C. jejuni*, although there have been few studies that examine the role of signal transduction in biofilm formation. It has been suggested that the impaired motility of aflagellated mutants impacts the access of bacterial cells to an abiotic surface [18,19]. Mutations to *cheW* and *cheV* proteins also serve to diminish motility in *C. jejuni*, as demonstrated by the decreased migration in Fig. 4. However, despite the decrease in motility, both *ΔcheW* and *ΔcheV* exhibited stark increases in the amount of biofilm formed, consistent with the significant biofilm-associated down-regulation of chemotaxis and chemosensory genes / proteins as determined by omics analysis. We propose that this *che*-associated reduced motility may provide more time for planktonic cells to adhere, also resulting in our observed increase in autoagglutination. This would allow more efficient formation of microcolonies during the initial stages of biofilm formation in *C. jejuni* (S3 Fig) and, thereby, a higher biomass is mature biofilms. All these data combine to suggest that regulation of signal transduction in *C. jejuni* may be critical during biofilm formation.

*C. jejuni* biofilms play a functional role in the ability of this organism to withstand environmental stresses and, thereby, impact the ability to infect human hosts. Whilst chemotaxis has been suggested to play an important role, motility has been the main focus in regards to biofilm formation. We have provided evidence that diminished motility is conducive to biofilm formation in *C. jejuni*, and that chemotaxis may play a larger role than previously thought. This presents the possibility that biofilm formation in *C. jejuni* may be regulated through modulation of chemotaxis.

## Materials and methods

### Bacterial strains

The wild-type *C. jejuni* strain NCTC 11168 O used in this study was provided by Dr. Diane Newell [21]. *ΔcheV* and *ΔcheW* mutant strains were cloned into 11168 O using plasmid constructs provided by Professor Julian Ketley, University of Leicester, United Kingdom as described previously [16]. All strains were grown microaerobically for a period of 12 hrs at 42°C on Columbia Blood Agar (CBA) with Skirrow supplement, and where appropriate, supplemented with 20 µg/mL Chloramphenicol.

Planktonic cultures of *C. jejuni* strains for use in proteomic and transcriptomic analysis were grown from an overnight plate culture, which was used to inoculate 250 mL of heart infusion (HI) broth (Oxoid, United Kingdom). These cultures were incubated under microaerobic conditions at 42°C and 120 x *rpm* for 12 hrs. For biofilm growth, an overnight culture of *C. jejuni* was diluted to an OD600 of 0.75 in Mueller Hinton (MH) broth (Oxoid). 10 mL of the cell suspension was then inoculated into glass petri dishes and incubated aerobically at 42°C.

### RNA sequencing

Planktonic and biofilm samples of *C. jejuni* were prepared for transcriptomic analysis as described [18].

Cells were harvested in 4 M guanidine thiocyanate (Promega, United States) in a 1% sodium lauryl sarcosine (Sigma, United States) solution. Samples were carefully laid over a 3.5 mL cushion of 5.7 M cesium chloride (Sigma) in 0.1 M EDTA (ChemSupply, Australia) in an OptiSeal polypropylene centrifuge tube (Beckman Coulter, United States) with the remaining volume of the tube being filled with Diethyl pyrocarbonate (DEPC) treated water. Samples were centrifuged at 27,000 x *rpm* for 16 hrs at 4°C in an Optima L-100 XP ultracentrifuge (Beckman Coulter) using a SW32Ti rotor.

Following centrifugation, the upper phase was removed and the tubes were cut before being allowed to completely drain. The RNA pellet was washed in 100 µL 70% ethanol before being resuspended in 120 µL Tris EDTA (TE) / Sodium dodecyl sulfate (SDS) with 80 µL of TE being used to dissolve remaining RNA. 650 µL of chilled ethanol was added to resuspended samples in addition to 10 µL sodium acetate (Sigma) to precipitate RNA. Samples were centrifuged at 14,000 x *rpm* in a Microfuge 22R centrifuge (Beckman Coulter) for 15 mins at 4°C. The supernatant was removed and a further 650 µL of 75% ethanol was used to wash the pellet. Samples were centrifuged for a further 5 mins at 14,000 x *rpm* and the supernatant removed. Pellets were allowed to air dry before resuspension in RNase free water. Quality and concentration of RNA was assessed using a Nanodrop 2000 (Thermo Scientific, United States).

RNA sequencing analysis was conducted as detailed [18].

### Preparation of peptide samples for analysis by LC-MS/MS

*C. jejuni* samples for use in proteomic analysis were grown in a similar manner as those used in RNA sequencing. Following incubation, samples were harvested in RNase free water. Frozen, washed bacterial cell pellets were lyophilised overnight and kept at -80°C until required. Pellets were resuspended in lysis buffer containing 150 mM Tris-HCl, 125 mM NaCl and 0.1 mm acid-washed glass beads (Sigma) and lysed by 4 rounds of bead-beating (4 m/s, 1 min) with 1 min rest periods on ice. Cell debris was removed by centrifugation at 16,000 x *g* for 15 mins at 4°C. 250 µL of sample was mixed with ice-cold water/methanol/chloroform in a ratio of 3:4:1 to precipitate proteins. Proteins were resuspended in 6 M urea, 2 M thiourea and reduced with dithiothreitol (DTT; 10 mM) at 37°C for 1 hr followed by alkylation with iodoacetamide (IAA; 20 mM) for 1 hr at room temperature in the dark. Samples were then diluted 10-fold in 100 mM triethylammoniumbicarbonate (TEAB) and quantified using the Qubit protocol (Life Technologies, Carlsbad CA). Samples were digested with trypsin in a ratio of 1:50 enzyme/sample for 16 hrs at 37°C. Lipids were precipitated using formic acid (FA) and removed by centrifugation at 16,000 x *g* for 15 mins at 4°C.

Supernatants were acidified with trifluoroacetic acid (TFA) to a final concentration of 0.1%, and peptide purification was performed using 60 cm^3^ hydrophilic lipophilic balance (HLB) cartridges (Waters Corp., Bedford MA). Cartridges were activated with 100% methanol (1 volume), followed by 100% acetonitrile (MeCN; 1 volume) and 70% MeCN / 0.1% TFA (1 volume). The cartridges were equilibrated with 0.1% TFA (2 volumes) and loaded with peptide sample. Samples were reapplied three times to ensure sufficient binding, washed with 0.1% TFA, and eluted with 70% acetonitrile / 0.1% TFA (1 volume). Peptides were then lyophilised and resuspended in 100 mM TEAB and quantified using Qubit. 125 µg of peptides were labelled per channel using isobaric tags for relative and absolute quantitation (iTRAQ) according to the manufacturer’s protocol (SCIEX, Framingham, MA). Samples were combined and diluted to 1 mL in 0.1% TFA and purified by HLB columns, as described above. Labelled samples were lyophilized and stored at -80°C until required.

### Quantitative LC-MS/MS of peptides from *C. jejuni*

8 µg of iTRAQ-labelled peptides were separated into 10 fractions by offline hydrophilic interaction liquid chromatography (HILIC) using an Agilent 1100 chromatography system. Fractionation was performed using a 20 cm, 320 µm inner diameter (i.d) column packed with TSK-Amide 80 HILIC resin, 3 µm particle size (Tosoh Biosciences, Tokyo, Japan). Samples were resuspended in buffer B (90% MeCN / 0.1% TFA) and separated using a linear gradient: sample loading for 10 mins with 100% buffer B at 12 µL / min, sample elution from 90 - 60% buffer B at 6 µL/min for 40 mins. Peptide elution was monitored by an absorbance detector at 280 ± 4 nm. Fractionated samples were lyophilised and stored at -20°C until mass spectrometric analysis.

HILIC fractionated peptides were resuspended in 0.1% FA and separated on an Ekspert NanoLC 400 system coupled to a SCIEX TripleTOF 6600 quadrupole time-of-flight mass spectrometer (SCIEX, Framingham MA) operated using AnalystTF (v.1.7.1). Peptides were loaded in buffer A (0.1% FA) directly onto an in-house packed 75 µm x 55 cm reversed phase column (1.9 µm particle size, C18AQ; Dr Maisch, Germany), then separated by adjusting the proportion of buffer B (80% MeCN, 0.1% FA) from 5–40% over 120 mins at 400 nL/min at 55°C. The TripleTOF 6600 was operated in positive ion mode using information (data) dependant acquisition mode, with the top 40 most intense ions with a minimum ion count of 400 and charge state between +2 and +5 selected for MS/MS fragmentation. MS scans were acquired using a mass range of 350 –1400 *m/z* and an accumulation time of 250 ms, with the instrument operated in high sensitivity mode using a mass range of 100-1800 *m/z*, accumulation time of 50 ms, resolution set to unit, mass tolerance set to 20 ppm and the options for rolling collision energy (CE) and adjust CE for iTRAQ reagent selected.

LC-MS/MS data were identified and quantified with ProteinPilot (5.0.0.0) using the Paragon algorithm, searched against the UniProt *C. jejuni* NCTC 11168 genome database (UP000000799; organism ID 192222; release May 24, 2018, 1623 proteins) with search parameters as follows: sample type set as iTRAQ 4-plex (Peptide Labeled), cysteine alkylation with IAA, digestion using trypsin allowing one missed cleavage, instrument set as TripleTOF 6600, search effort as thorough and FDR analysis using a detected protein threshold of 0.05. Total reporter ion intensity for each protein was calculated at the protein level, normalised against the summed reporter ion intensity for all proteins including contaminants present in that channel. An average of reporter ion intensity at the protein level was used for downstream analysis. Proteins were included for analysis only if they contained ≥2 identified and quantified peptides. A difference of ≥ +/-1.5 fold was considered differentially abundant between groups.

### Post-processing of transcriptomics and proteomics data

Heat map data were generated in Morpheus (https://software.broadinstitute.org/morpheus/). Functional cluster analysis was performed in STRINGdb [22].

### Assessment of biofilm formation

Plate-grown cultures were harvested with 1 mL of MH Broth and the OD600 of samples was adjusted to 0.75. 100 µL of bacterial suspension was used to inoculate a 96-well plate, which was then incubated for 48 hrs at 42°C. Wells were gently washed with 200 µL of Phosphate Buffered Saline (PBS) and stained with 100 µL 1% Crystal Violet (CV) for 15 mins. Wells were gently washed a further 3 times with 200 µL of PBS and biofilm solubilised with 100 µL of biofilm solvent as previously published [23]. Plates were then analyzed using a Victor X multilabel plate reader at 600nm. Statistical significance was determined using a paired, two-tailed students t-test.

### Motility assays

Motility assays were conducted as previously described [14]. Following incubation, cells were harvested in 1 mL MHB and OD600 adjusted to 0.5. 5 µL of this bacterial suspension was then stabbed into 0.35% MH plates and incubated for 48 hrs at 42°C. The diameter of the measured growth from the stab point was measured in mm. All motility assays were performed in triplicate.

### Time lapse microscopy

Cells were harvested from plates in 1 mL of MHB and adjusted to an OD600 of 0.75. Bacterial suspensions were used to inoculate 24-well plates containing sterilised glass cover slips. Plates were incubated at 42°C for 45 hrs. At each five-hour timepoint, cover slips were gently washed with PBS and fixed with 5% formalin. Cover slips were then gently washed with 1 mL of PBS and stained with 500 µL of CV for 15 mins. CV was then removed and the cover slips were washed three times in PBS before being allowed to air dry and mounted to slides. Samples were analysed using a Nikon Eclipse E600 light microscope and images taken using a Nikon DXM-1200C digital camera and Act-1 visualisation software.

### Autoagglutination assays

Autoagglutination assays were carried out as previously described [14]. Cells were harvested from incubated plates in 1mL of PBS and suspensions were diluted to an OD600 of 1.0. 2mL of the diluted bacterial suspensions were inoculated into 10mL glass culture tubes and incubated for a period of 24 hours at 42°C. The culture tubes were then inspected for agglutination of cells and the upper 1mL was assessed spectrophotometrically at 600nm.

## Acknowledgements

The authors would like to thank Professor Julian Ketley for providing the CheV and CheW mutant strains and Dr Dianne Newell for providing access to the *C. jejuni* strain 11168 O.

## Supporting information captions

**Fig S1a. STRINGdb cluster analysis**. DE genes with fold-change >+2, using custom assigned confidence of 0.850 and orphan nodes removed. 318 nodes, 456 edges with PPI enrichment p-value <1.0e-16

**Fig S1b. STRINGdb cluster analysis.** DE genes with fold-change >+5 (118 genes), using custom assigned confidence of 0.850 and orphan nodes removed. 118 nodes, 96 edges with PPI enrichment p-value <1.0e-16

**Fig S1c. STRINGdb cluster analysis.** DE genes with fold-change <-2, using custom assigned confidence of 0.850 and orphan nodes removed. 301 nodes, 283 edges with PPI enrichment p-value <2.2e-16

**Fig S1d. STRINGdb cluster analysis.** DE genes with fold-change <-5 (96 genes), using custom assigned confidence of 0.850 and orphan nodes removed. 97 nodes, 23 edges with PPI enrichment p-value = 0.00043

**Fig S2a. STRINGdb cluster analysis.** DA proteins with fold-change >+1.5, using custom assigned confidence of 0.850 and orphan nodes removed. 220 nodes, 314 edges with PPI enrichment p-value <2.6e-11

**Fig S2b. STRINGdb cluster analysis.** DA proteins with fold-change >-1.5, using custom assigned confidence of 0.850 and orphan nodes removed. 210 nodes, 173 edges with PPI enrichment p-value <2.21e-10

**Fig S3. Time lapse microscopy.** Demonstrating the tendency for aggregation of 11168-O, Δ*cheV* and Δ*cheW* isogenic mutants. Both *ΔcheV* and *ΔcheW* strains demonstrated a significantly higher tendency to cluster into microcolonies when compared to wild type 11168-O over a period of 25 hours.

**Table S1. RNAseq complete.**

**Table S2. RNAseq DE genes.**

**Table S3. Proteomics complete.**

**Table S4. Proteomics DA proteins.**

**Table S5. Comparative RNAseq and proteomics.**

